# Diversity loss in microbial ecosystems undergoing gradual environmental changes

**DOI:** 10.1101/2023.07.07.548183

**Authors:** Aviad Berger, Maya Gatt Harari, Avner Gross, Amir Erez

## Abstract

Microbial ecosystems in soils, oceans, and other environments are essential for global ecological stability. Environmental shifts are anticipated to trigger destabilizing events across the planet. In this study, we model how gradual environmental changes impact ecosystems, specifically when leading to a loss of diversity. We investigate how an ecosystem, within a serial-dilution setup, relaxes to a stable steady state. Our results reveal that as an ecosystem approaches its loss of diversity transition, its dynamics slow down. Consequently, diverse ecosystems, such as tropical rainforest soils, gradually driven past their transition point may exhibit a significant response lag. This suggests that some ecosystems may be closer to a collapse in diversity than current observations indicate. Although our model does not capture the full complexity of real-world ecosystems, it highlights critical aspects underlying the loss of biodiversity in changing environments. This has potential implications for empirical studies and when planning interventions.

## INTRODUCTION

Microbial communities pervade our planet as essential constituents in every ecosystem, ranging from human guts to rainforest soil and coral reefs. These omnipresent microbes are engaged in a relentless struggle for finite resources, a competition possibly as ancient as life itself. Alongside competition, microbial communities are defined by their remarkable diversity, supporting the coexistence of thousands of species. How this diversity perseveres, despite the aforementioned competition, remains a central question (Chesson, 2000; Hu, Amor, Barbier, Bunin, & Gore, 2022; Rubin, Ispolatov, & Doebeli, 2023). However, slow yet cumulative environmental changes can provoke a collapse in diversity, leaving ecosystems populated by only a handful of surviving species. We sought a minimal description of how diversity is affected in response to the slow dynamics of environmental changes.

A recent global investigation highlighted that the carbon use efficiency (CUE) of soil microbiota is a crucial factor governing global soil organic carbon storage and fluxes (Tao et al., 2023). Notably, the CUE is lowest in tropical rainforest soils, contributing to their low carbon stocks and rapid turnover. Furthermore, a correlation exists between CUE and microbial diversity (Domeignoz-Horta et al., 2020). With these insights, we have chosen microbes in tropical rainforest soils as a specific exemplar of the diversity transition underpinning this study. We consider a plausible scenario of a future collapse in the diversity of microbes within these soils.

Species diversity, which comprises the quantity and relative abundance of species within an ecosystem, is intertwined with ecosystem health and resilience. Ecosystems can destabilize if their diversity diminishes due to external pressures, resulting in the loss of essential functions and heightened susceptibility to disturbances (Veraart et al., 2012; Scheffer, Carpenter, Dakos, & van Nes, 2015; Wu et al., 2022). Such functional loss in rain-forest soils could have catastrophic consequences, including the potential release of vast amounts of CO_2_ into the atmosphere.

The age and extreme weathering of rainforest soils often dictate their microbial processes and community structure via phosphorous availability, acting as a Leibig-like limiting nutrient (Cleveland & Townsend, 2006; Gross, Pett-Ridge, & Silver, 2018; Fleischer et al., 2019; Du et al., 2020; Oliverio et al., 2020; Hou et al., 2020). Climate change is projected to influence microbial phosphorous limitation in two main ways: firstly, by diminishing the bio-availability of soil inorganic phosphorous through altered rainfall patterns affecting redox-sensitive reactions, which govern the release and re-absorption of phosphorous from soil minerals; secondly, by inducing phosphorous dilution in plant organic matter due to anthropogenic CO_2_ fertilization (Loladze, 2002; O’Connell, Ruan, & Silver, 2018; Gross, Lin, Weber, Pett-Ridge, & Silver, 2020). Collectively, these transformations suggest the possibility of rainforest soil microbial ecosystems approaching critical transition points, with significant implications for the global carbon cycle.

It has been known for decades that varied physical phenomena share the same scaling behavior near a transition point (Goldenfeld, 1992). In recent years, efforts were made to include in such analyses biological systems (Mora, Walczak, Bialek, & Callan, 2010; Muñoz, 2018; Erez, Byrd, Vogel, Altan-Bonnet, & Mugler, 2019; Byrd et al., 2019; Bose & Ghosh, 2019; Erez, Byrd, Vennettilli, & Mugler, 2020; Vennettilli, Erez, & Mugler, 2020) and ecological systems (Dai, Vorselen, Korolev, & Gore, 2012; Dakos & Bascompte, 2014; Biroli, Bunin, & Cammarota, 2018; Hu et al., 2022). Near a transition, slowing down of dynamics typically occurs, so response to a change takes a long time to settle. Due to this long, emergent time scale, short-scale differences in systems can average out. Thus, many systems can be described by the same scaling relations between observable quantities. Such scaling has been influential in elucidating various biological phenomena (Tkačik & Bialek, 2016; Bradde & Bialek, 2017; Biroli et al., 2018; Bose & Ghosh, 2019).

In the field of ecology, there has been extensive discussion on early warning indicators of ecosystem collapse and the slowing-down of dynamics (Gandhi, Levin, & Orszag, 1998; Dai et al., 2012; Dai, Korolev, & Gore, 2013; Dakos & Bascompte, 2014; Scheffer et al., 2015). Recent advancements include a deep-learning method detecting early-warning signs of tipping points (Bury et al., 2021); using ‘exit time’, the interval between switching events, to quantify resilience in lake cyanobacteria and other systems (Arani, Carpenter, Lahti, van Nes, & Scheffer, 2021); other early warning signals for critical transitions in ecological systems (Xu, Patterson, Levin, & Wang, 2023). All these approaches crucially involve emergent timescales in describing ecological transitions. While various authors have proposed distinct measures of ecological stability, one might agree stability should encapsulate elements of biodiversity, resilience, and resistance (Van Meerbeek, Jucker, & Svenning, 2021). With these foundations in mind, we asked how emergent critical slowing down manifests in the species diversity in a changing environment.

In this manuscript, we focus on consumer-resource models in a serial dilution framework, an under-explored theoretical framework. We consider an ecosystem’s diversity as it is slowly driven across its loss-of-diversity transition point. In the context of rainforest soils, our theory may pertain to changes in the concentration of two forms of phosphorous, a limiting nutrient, in relation to its redox-dependent bio-availability. This driving action results in a delayed response, with the lag more pronounced in ecosystems boasting a higher species richness. Although suggestive of the concept of ‘extinction debt’ (Tilman, May, Lehman, & Nowak, 1994), the lag we observe stems from the slowing down of dynamics near a transition. Given the ubiquity of critical slowing down across various transitions, it is plausible that our analysis extends beyond the specific model and ecological example we examined.

## MATERIALS AND METHODS

### Model

To model microbial dynamics, we used a serial dilution framework in which *m* species compete for *p* nutrients within a series of recurring batches (Erez, Lopez, Weiner, Meir, & Wingreen, 2020). In alignment with Leibig’s law, we posit these nutrients as interchangeable forms of a solitary limiting resource, such as carbon or phosphorous. We concentrate on a case involving two forms of this nutrient. At the commencement of a batch, an initial nutrient composition is provided, 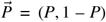, such that the total nutrient concentration, 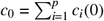, and *c*_*i*_(0) = *c*_0_ *P*_*i*_ is the concentration of nutrient *i* at time *t* = 0. The nutrient composition can be plotted on a simplex (Fig. 1a). Time is measured from the beginning of the batch. Concurrently with nutrient provision, a batch is inoculated with *m* different species such that 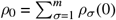, where *ρ*_*σ*_(*t*) represents the biomass concentration of species *σ* at time *t*. In the numerical calculations in this manuscript, we used maximum-entropy initial conditions, 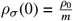, reflecting minimal prior knowledge about the system (J. Jost, 2020).

**FIGURE 1.**
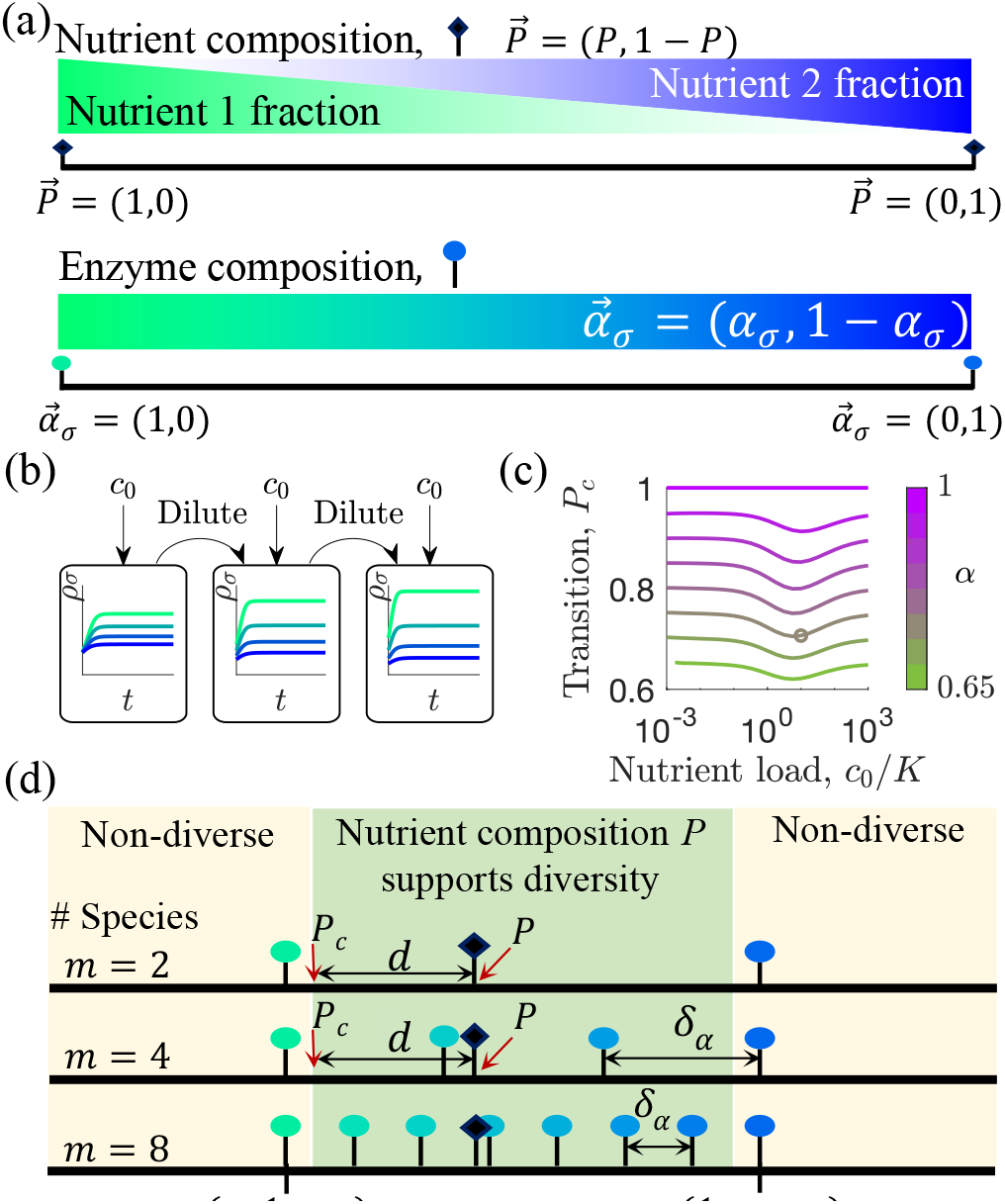
Schematic of the model. (a) Representation of the nutrient supply composition *P* = *c*_1_(0)/*c*_0_ (black diamond) on a 2-nutrient simplex, where the left and right endpoints correspond to *P* = 1 and 0, respectively. Representation of the metabolic strategies, *α*_*σ*_, (colored circles) on the same simplex. (b) Illustration of the relaxation of the serial dilution process. Each cycle of batch growth begins with cellular biomass densities *ρ*_*σ*_ and total nutrient concentration *c*_0_. The system evolves according to Eqs. 1-2 until nutrients are completely consumed. A fraction of the total cellular biomass is then used to inoculate the next batch again, repeating until steady state. (c) The fraction of Nutrient 1 at the transition point, *P*_*c*_, depends on the total amount of nutrient provided. Colors correspond to different systems, each with 2 species with *α*_*σ*_ = (*α*, 1 – *α*) and (1 – *α, α*). The circle represents the *c*_0_ and α used in panels c-d and in Figures 2-4. (d) Here, the nutrient supply (black diamond) is inside the region that supports multi-species coexistence (see text). This defines a distance, from the nutrient supply to the edge of the coexistence region, *d* = *P*_*c*_ – *P*, where *d* = 0 marks the transition point and *d* > 0 is the coexistence state. Three systems are shown with *m*=2,4,8 equally spaces strategies. Inoculating with *m* species with symmetric, equally spaced strategies bounded by *α* and (1 – *α*) defines a distance between two neighboring strategies, 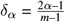.

How is nutrient consumption modeled in our framework? Each species is allocated a fixed and immutable ‘metabolic enzyme budget’, employed to consume one or multiple nutrient types—its metabolic strategy (Posfai, Taillefumier, & Wingreen, 2017; Erez, Lopez, et al., 2020; Erez, Lopez, Meir, & Wingreen, 2021). We index a species by *σ*, defining it by this strategy, 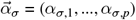, where *α*_*σ,i*_ denotes the metabolic capacity of nutrient *i*, i.e., its maximum uptake rate. We impose a tradeoff, that every species maintains an identical metabolic budget, ∀*σ*: Σ_*i*_ *α*_*σ,i*_ = *E*_*σ*_ = 1, wherein assigning the value 1 defines the units in which time is measured (Posfai et al., 2017). Consequently, the metabolic enzyme strategy can be placed on the same simplex as the nutrient composition (Fig. 1a). The uptake rate of nutrient *i* also depends on its concentration, *c*_*i*_(*t*), following a Monod form (Monod, 1942), given by 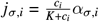, where *K* signifies the half-saturation constant, which we assume is uniform across all nutrients for the sake of simplicity. A more general form of the model is described in (Erez, Lopez, et al., 2020). The dynamics of the population and nutrients within a batch conform to (MacArthur, 1970; Chesson, 1990; Erez, Lopez, et al., 2020),

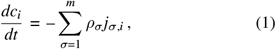

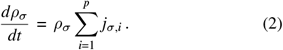

The serial-dilution batch-culture setup, despite its widespread experimental use and real-world analogies, is theoretically under-explored. Once all nutrients within a batch are depleted, a new batch is started. This new batch, introduced with the same initial nutrient concentration and an *inoculum* comprising microbes with a total concentration of *ρ*_0_, is started. In the serial dilution setup, the inoculum is composed of the species composition at the end of the previous batch, post dilution (Fig. 1b). In essence, a batch is inoculated with microbes and nutrients. After the nutrients are consumed, the final composition is diluted and used to inoculate the next batch, thus continuing the cycle.

For these equations to reach a steady state, the relative abundances of species at the beginning and end of the batch must be identical. In other words, all surviving species must exhibit equivalent growth (as gauged by fold change) within a batch. Any disparity in growth between species could lead to competitive exclusion, where dominant species outcompete the others. In our previous works (Erez, Lopez, et al., 2020; Erez et al., 2021), we analyzed the steady states of this serial dilution setup and identified the conditions required for stable diversity. At low nutrient concentrations (*c*_0_ ≪ *K*), the serial dilution setup mirrors the behavior of a chemostat, a system where nutrients are continuously supplied and diluted (Posfai et al., 2017). The chemostat limit supports an unlimited number of coexisting species, subject to a graphical condition: the nutrient composition must be enveloped by the convex hull of metabolic strategies, which are bounded by the extreme strategies on the simplex (Posfai et al., 2017). We observed that when increasing *c*_0_ beyond the chemostat limit, so that *c*_0_ ≈ *K*, the convex hull rule still holds true, albeit with ‘remapped’ boundaries that delimit the range of nutrient compositions capable of sustaining coexistence. The remapping changes the transition point, from the most extreme strategy, to *P*_*c*_, and depends on *c*_0_ as shown in Fig. 1c. When the nutrient supply composition is within this remapped region, stable multi-species coexistence is possible. An example of a diverse community where the nutrient supply *P* is nestled within the coexistence region demarcated by *P*_*c*_ is shown in Fig. 1d. This defines a distance to the loss-of-diversity transition point, *d* = *P*_*c*_ – *P*, where *d* > 0 signifies the region of the coexistence steady state.

### Relaxation to steady state

How long does it take the system to relax to steady state? How does this ‘relaxation length’ depend on the proximity to the loss-of-diversity transition point, *d* → 0? The relaxation length quantifies the system’s responsiveness to changes, an indicator of ecological resilience (Scheffer et al., 2015). For example, in serial-dilution cultures of *Saccharomyces cerevisiae*, the number of batches required to revert to steady state following a salt shock diverges near the system’s tipping point (Dai et al., 2012). This number of batches, emerging from the system’s collective dynamics, is what we term the relaxation length, *ξ*.

When the relaxation length grows, the system takes longer to respond to change and settle, showing *critical slowing down*. The physics literature is replete with examples of critical slowing down near transition points (Goldenfeld, 1992), and the ecological literature has many examples where slowing down serves as a precursor to collapse (Gandhi et al., 1998; Dai et al., 2012, 2013; Dakos & Bascompte, 2014; Biroli et al., 2018; Scheffer et al., 2015; O’Keeffe & Wieczorek, 2020).

To extract the relaxation length from simulations, we note that near the steady state, convergence is dominated by a single timescale. Consequently, an observable measure such as the Shannon diversity approaches steady state as *S*_*b*_ – *S*_∞_ ∼ *e*^−*b*/*ξ*^, with *b* representing the batch number, the Shannon diversity defined as 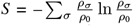, and the steady state diversity given by *S*_∞_. To stay close to steady state, we consider only values of *b* such that (*S*_*b*_ – *S*_∞_) <(*S*_0_ – *S*_∞_)/100. As such, *ξ* should not depend on initial conditions near steady state—a fact that we have numerically confirmed. We extracted *ξ* by fitting the exponential form, as shown in Fig. 2a (more details are in the *Supplemental Material* Sec. A).

**FIGURE 2.**
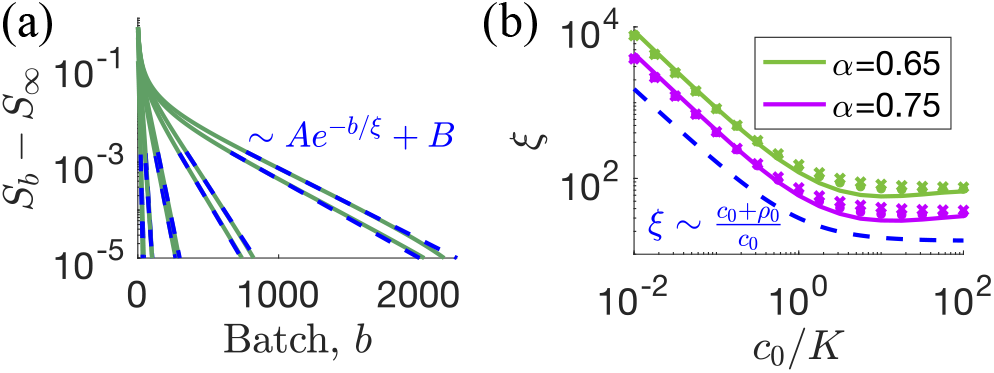
Relaxation to steady state. (a) The convergence of the Shannon diversity to steady state, for various distances from the transition point (green curves). The fit to *S*_*b*_ – *S*_∞_ ∼ *Ae*^−*b*/*ξ*^ + *B* (dashed blue) is imposed on the numerical results, allowing us to extract *ξ* (*cf*. Supplemental Material Sec. A). (b) How *ξ* depends on *c*_0_ (curves), for systems with two species, (*α*, 1 – *α*) and (1 – *α, α*), and *α* = 0.65, 0.75. The dependence on nutrient amount, 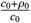, is shown in dashed blue. Also shown as dots and crosses, the approximate form, *ξ*_*chem*_ according to Eq. 4 and Eq. 5, respectively. The distance to the transition point, *d* ≡ *P*_*c*_ – *P* = 10^−2^, where *P*_*c*_ depends on *c*_0_ according to Fig. 1c.

In order to exclude potential artifacts that could be introduced by the Runge-Kutta method used to integrate the dynamics, we also took into account an alternative stochastic version of the model. To compute the correlation length (defined in batches) of the stochastic dynamics, we implemented Gillespie simulations (Erez et al., 2021) (*Supplemental Material* Sec. B). In the stochastic version, we first let the system reach steady state, and then measured the auto-correlation of the stochastic fluctuations. Conceptually, the stochastic system can be thought of as a vibrating ‘marble in a cup’ with the auto-correlation time pertaining to the marble’s location (Arani et al., 2021). As expected, the correlation time of the stochastic dynamics is equivalent to the relaxation length of the deterministic dynamics, demonstrated in Fig. S1. Henceforth, we only consider the relaxation of the deterministic dynamics.

## RESULTS

### Analytical form of the relaxation length

To derive an approximate form for the relaxation length we consider the case where the initial nutrient load is small, *c*_0_ ≪ *K*.

In this case, the steady state for the serial dilution is the same as the steady state where nutrients are continuously supplied, which is known as a chemostat (Erez, Lopez, et al., 2020). We used a first-order expansion of the system at small *c*_0_ to calculate the chemostat-limit dependence of the relaxation length, *ξ*_*chem*_, close to the loss-of-diversity transition point, where *d* ≪ 1. For the full derivation, refer to the *Supplemental Material* Sec. C. For a situation where there are two species with strategies (*α*, 1 – *α*) and (1 – *α, α*), and two nutrients such that *c*_1_(0) + *c*_2_(0) = *c*_0_, at the chemostat limit (*c*_0_ ≪ *K*), the relaxation length *ξ*_*chem*_ follows:

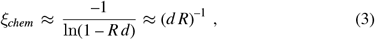

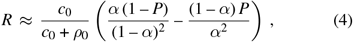

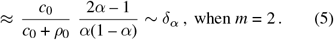

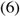

Here, 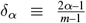 is the distance between neighboring strategies (Figure 1d). A comparison between Eq. 3 and the numerical value of *ξ* reveals a striking agreement even for *c*_0_ ≫ *K* (Fig. 2b), capturing the α dependence well. Moreover, even the rough approximation behind Eq. 5 which takes *P* ≈ *α*, works well. In the derived equation, we introduced an explicit dependence on *c*_0_ as 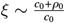 because when *c*_0_ ≪ *ρ*_0_, a minimum of *ρ*_0_/*c*_0_ batches are needed for growth. The difference between the numerically calculated and approximate values when *c*_0_/*K* ≈ 10 can be attributed to the faster convergence induced by the ‘early-bird’ effect (Erez, Lopez, et al., 2020). The decoupling of *d* and *c*_0_ in the scaling of *ξ* allows us to focus the rest of this paper on *c*_0_/*K* = 10. We have also verified numerically that the following results apply to other values of *c*_0_.

### Slower relaxation in multi-species ecosystems

In ecosystems composed of many species, a crucial question is how the relaxation length, or the time required for a system to return to equilibrium following a disturbance, changes with species richness. We simulated systems with varying species counts—*m* = 2, 4, 8, …, 1024—and determined their respective relaxation lengths. We found that the divergence of *ξ* as *d* → 0 is different for *d* < 0 and *d* > 0 (Fig. 3a). Plotted on a logarithmic scale, the difference in behavior is even more striking (Fig. S2a). When *d* < 0, the steady state cannot support diversity. As such, when the system is near the steady state, it effectively becomes a two-species system. Consequently, for all *m* values, *ξ* ∼(–*d*)^−1^ (Fig. 3b) with a full scaling form according to Eq. 5, *ξ* ∼(–*d δ*_*α*_)^−1^ (Fig. S2b).

**FIGURE 3.**
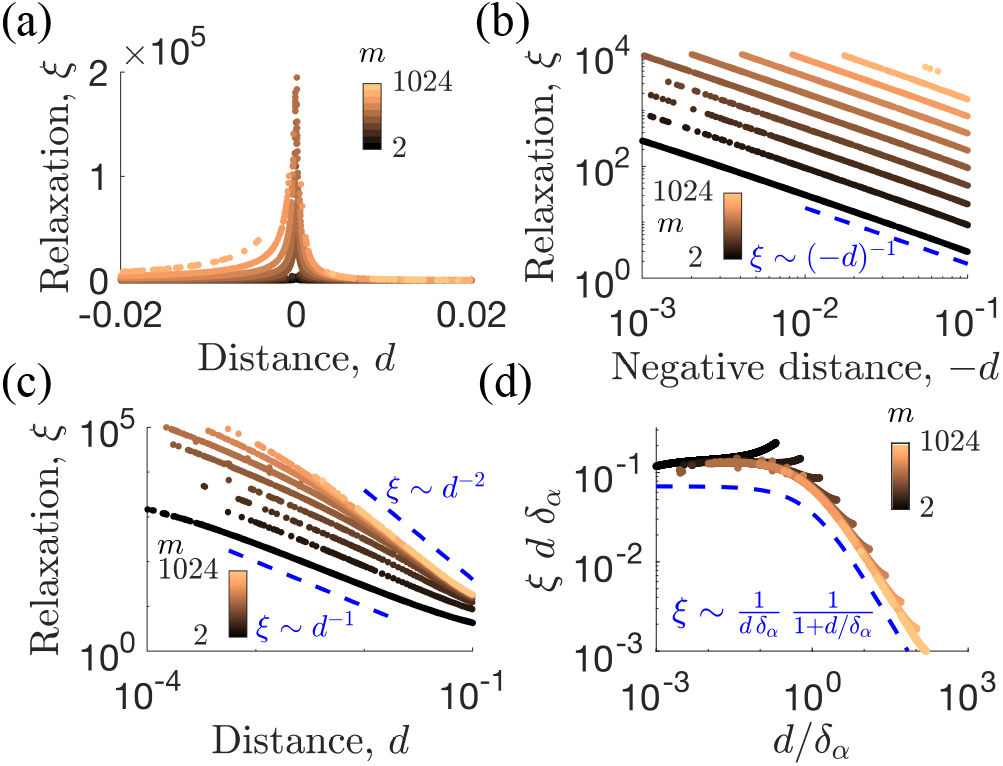
The number of batches required for relaxation, *ξ*, as a function of the distance to the transition point, *d*, with *d* > 0 marking the diverse steady state; *m*=2,4,…,1024 species (colors) initially inoculated in equal amounts with metabolic strategies equally spaced between (*α*, 1 – *α*) and (1 – *α, α*), and *α*=0.75, *c*_0_=10, *K*=*ρ*_0_=1. (a) *ξ* for different *m* values diverges at *d*=0 but is not symmetric for positive and negative *d*. (b) On a logarithmic scale, where *d* < 0 and the nutrient composition does not support coexistence. Relaxation to a single species occurs with *ξ* ∼(–*dδ*_α_)^−1^ (see Eq. 3). (c) For positive *d*, the log-scale reveals a non-trivial dependence of *ξ*(*d*) on the number of species. (d) Scaling collapse for 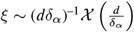. Dashed blue: the Ansatz 𝒳(*x*) = (1 + *x*)^−1^.

At the coexistence state, *d* > 0, the scaling of *ξ* with *d* is more complicated. A logarithmic plot reveals that, although for *m* = 2, as in Eq. 3, *ξ* ∼ *d*^−1^, a crossover to *ξ* ∼ *d*^−2^ behavior appears at larger *m* values (Fig. 3c). This crossover depends not on the absolute distance to the transition point, but rather on the relation between the absolute distance *d* and the smallest distance in enzyme allocation between species, given by 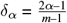. A scaling hypothesis, *ξ* ∼(*dδ*_α_)^−1 𝒳^ (*d*/*δ*_α_) with an unknown universal function 𝒳(*x*), successfully collapses the data (Fig. 3d). Furthermore, a form 𝒳(*x*) = (1 + *x*)^−1^ seems to capture the results effectively. Therefore, more species-rich systems (*m* ≫ 2 → *δ*_α_ ≪ 1) significantly slow down the relaxation dynamics near the transition point compared to smaller systems, transitioning from *ξ* ∼ *d*^−1^ to *ξ* ∼ *d*^−2^. The dependence on *d*/*δ*_α_ suggests that the degree of slowdown is determined by the number of species that remain (when visualized on the simplex) between the nutrient supply *P* and the transition point *P*_*c*_. These observations are applicable for other values of *c*_0_ (see, e.g., Fig. S3). In summary, the more species in the ecosystem, the slower the system responds near the loss-of-diversity transition.

### Lagged ecosystem response to a gradual change

How does the slowing-down presented influence our understanding of large ecosystems? Let us consider tropical rainforest soil, where phosphorous is a limiting nutrient and comes in sub-stitutable forms that are metabolized differently (Gross et al., 2020). The changes in the availability of these forms, perhaps due to the effects of climate change on the soil environment, can be modeled as a slowly changing nutrient composition, crossing the loss of diversity transition point in the process. To represent the species diversity we used the *effective number of species, m*_*e*_ = *e*^*S*^, where the Shannon entropy 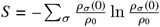 (L. Jost, 2006). Regardless of the number of species initially present, as the system approaches the transition point (*d* → 0), it eventually reaches a steady state where only a single species remains, which means *m*_*e*_ → 1 (as shown in Fig. 4a). Given uniform initial conditions and equally spaced metabolic strategies, it is possible to derive how *m*_*e*_ scales with the distance *d* from the transition point. At large *d*, when the nutrients are equally abundant, all species are equally abundant (Erez, Lopez, et al., 2020), therefore, *m*_*e*_ = *m* and 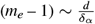 (Fig. 4b). At small *d* we have derived a logarithmic correction, 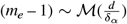, with ℳ(*x*) = *x*(1 – ln *x*). The correction to ℳ was derived using an Anzats, 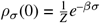 (*cf. Supplemental Material*, Sec. D).

**FIGURE 4.**
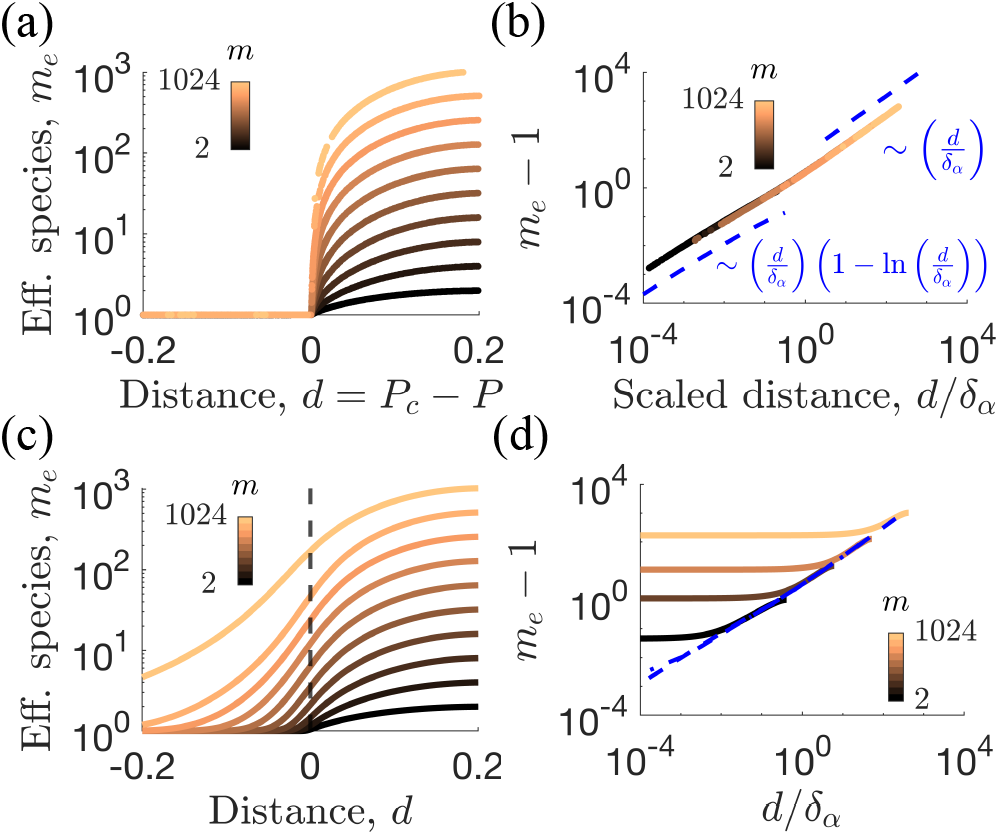
Species diversity quantified as the effective number of species, *m*_*e*_, vs. the distance to the transition point, *d*, for systems with *m*=2..1024 species (colors) initially inoculated in equal amounts with metabolic strategies *α*_*σ,i*_ equally spaced between (*α*, 1 – *α*) and (1 – *α, α*). Here, *K*=*ρ*_0_=1, *c*_0_=10, *α*=0.75. (a) At steady state, the effective number of species vs. *d*, showing how for *d* ≤ 0 only one species survives. (b) Scaling collapse onto a single curve which relates diversity to the distance to the loss of diversity transition point. Dashed blue: approximate forms according to Eq. S8. There is a small, but distinct slope difference between the large and small *d*/*δ*_*α*_ curves due to a logarithmic correction in the latter. (c-d) Response to a gradual change in nutrient composition from (0.5,0.5) to (1,0) which crosses the transition point over 10^4^ batches. (c) Diversity remains large, temporarily, even when *d* ≤ 0. Dashed: *d* = 0 line as a guide to the eye. (d) When scaled as in panel b, as *d*/*δ*_*α*_ → 0, diversity ‘freezes out’, as in the KZ effect, at a much larger value than its equivalent steady state. Dashed blue: the steady-state value according to the scaling collapse in panel b.

In a changing environment, can we say that the system is at steady state? We might consider a system to be at steady state, even when subjected to changing conditions, if it can respond quickly enough to the change, so the change seems slow in comparison. Such changes are sometime called *adiabatic*. However, a gradual change that takes the system across a tipping point behaves differently, and cannot be adiabatic. Due to critical slowing down, no matter how slowly the change happens, close to the transition the system is even slower, and cannot catch-up to the change. In the physics literature this lag in response is known as the *Kibble-Zurek* (KZ) effect, investigated in a myriad of systems (Kibble, 1976; Zurek, 1985; Biroli, Cugliandolo, & Sicilia, 2010; Chandran, Erez, Gubser, & Sondhi, 2012; Byrd et al., 2019). Instead of paralleling the change, the system drops out of steady-state, ‘freezing-out’ the observed diversity when nearing *d* = 0, causing a lag in response. Even after the ecosystem has crossed its tipping point, the lag in response can delay the collapse in diversity. There would linger transient species that are doomed to disappear, yet are left temporarily in the community.

This lag in response can be viewed as an ‘extinction debt’ (Tilman et al., 1994), though in contrast to the original concept, the lag here is introduced because of the crossing of a critical threshold and therefore the divergence of *ξ*. As we have demonstrated, *ξ* grows sharper as the number of species increases (Fig. 3). Therefore, we expect the KZ freeze-out to become more pronounced at multi-species ecosystems, i.e., large values of *m*. This phenomenon is precisely what is observed, where the diversity remains well above zero for *m* = 1024 even after driving the system through *d* = 0 (Figure 4c).

In logarithmic scale, the characteristic Kibble-Zurek freeze-out (Kibble, 1976; Zurek, 1985; Chandran et al., 2012) is evident in the plateauing of diversity as *d* decreases, indicating that the system cannot keep pace with changes in the nutrient composition. As the number of species increases, the diversity at which freeze-out (compared to the steady-state value) occurs also increases (Figure 4d).

## DISCUSSION

We studied a microbial ecosystem undergoing a serial-dilution, consumer-resource process, specifically as it approached a loss-of-diversity transition point. We used tropical rainforest soils, a diverse ecosystem that is limited by a single nutrient, to illustrate the relevance of our modeling to natural ecosystems.

### Relaxation to a diverse state is slower in species-rich ecosystems

We first considered a two-species system where we could derive the scaling of *ξ* if only a small amount of nutrient is given in each batch. Previously, we have shown that the steady state of this low-nutrient limit is equivalent to a chemostat wherein nutrients are continuously added and diluted (Erez, Lopez, et al., 2020). Here, in this low-nutrient limit we derived the number of batches required for the system to relax to steady state. The expression we derived for *ξ*_*chem*_ agrees with the *ξ* values obtained from numerical simulations. This agreement provides confidence in our numerical procedure and substantiates our scaling hypotheses. We introduced the scaling 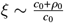 since if there is very little nutrient in the system then a single cycle of growth and dilution changes the population very little. Therefore, *ξ* scales with 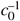 at small *c*_0_.

In serial-dilution cultures and absent in a chemostat, the ‘early-bird’ effect lets species gain an advantage through early growth. Despite being competitively inferior per capita in consuming residual nutrients after depleting their preferred type, early-bird species use their sheer numbers to starve their competitors (Erez, Lopez, et al., 2020). The early-bird effect has been recently confirmed in *ex-vivo* communities (Aranda-Díaz et al., 2023). Since the chemostat limit is equivalent to the serial dilution system at small *c*_0_, the difference between *ξ*_*chem*_ and *ξ* at intermediate nutrient amounts stems from higher-order terms—namely, the early-bird effect. The early-bird effect could be relevant to tropical rainforest soils since the limiting nutrient (P) often enters the system in pulses (Wood, Matthews, Vandecar, & Lawrence, 2016; Gross, Turner, Goren, Berry, & Angert, 2016; Reed et al., 2020).

For small enough *d*, in both the diverse steady state (*d* > 0) and non-diverse (*d* < 0) the relaxation follows the same *ξ* ∼(*d δ*_*α*_)^−1^. This is because near the transition, in both cases, the same single species survives and all the other species are out-competed. The relevant scaling of the distance to the transition point depends on the number of species as approximately 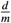 at large *m*. Therefore, using inocula that are very species-rich may result in slowing down of dynamics even when a less species-rich system may not be very close to the transition. This effect of the species richness of the inoculum on the relaxation to steady state may bear relevance to microbiome transplant applications, be it to humans or to crops.

Another effect emerges when relaxing to a diverse steady state: additional scaling with respect to *d*/*δ*_*α*_ which effectively changes the dynamics from *d*^−1^ to *d*^−2^. The correction to scaling is related to the number of species that remain beyond the guaranteed survivor, which can be expressed as 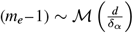. Intuitively, ℳ(*x*) ∼ *x* and although for small *x* there are logarithmic corrections to diversity, 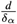 remains a relevant parameter combination. The number of species that exist between the nutrient composition and loss of diversity point influences the relaxation towards a diverse steady state, as indicated by the further dependence on 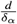. This dependence on the number of extant species may have implications for the relaxation dynamics of highly diverse ecosystems such as tropical soils—a species-rich ecosystem which is susceptible to environmental disruption (Bouskill et al., 2013; Lan et al., 2021).

### Response to gradual environmental change lags in multi-species ecosystems near collapse

A key finding of this study pertains to an environment undergoing a transition across the loss of diversity point, which we use as a minimal model for the effects of climate change. Due to critical slowing down, as the system approaches a transition, its response to change becomes slower, even when the change is slow. The observed diversity ‘freezes-out’, and lags in its response, resembling the Kibble-Zurek (KZ) effect in non-equilibrium physics (Kibble, 1976; Zurek, 1985; Byrd et al., 2019). The ecosystem’s collapse, even after passing its tipping point, is temporarily delayed due to the lag in its response. As higher diversity intensifies critical slowing down, the freeze-out should be more prominent in diverse ecosystems (Figure 3).

The KZ effect in response to climate change could be significant in natural ecosystems containing thousands of microbial species. Particularly concerning are tropical forest soils, where microbial diversity is known to depend on climatic and environmental factors (Zhou et al., 2016; Sun et al., 2020). Alarmingly, this would indicate that ecosystems that currently are recorded to have lost some, but not all of their diversity recently, may be well on the way to collapse, temporarily masked by the KZ lag. Because natural systems are spatially extended, this is likely to further slow down relaxation and generate even greater lag. Further relevance to natural ecosystems comes from two useful features of KZ-like sweeps: (i) knowledge of the precise transition point is not required, only that it is crossed at a constant rate; (ii) the precise temporal evolution of the parameters is irrelevant. Near the transition, the slowdown in dynamics is dominated by leading-order, linear behavior. This means that any change that crosses the transition point at a constant rate would result in a similar lag, regardless of its precise temporal dependence (Chandran et al., 2012). This makes KZ-like sweeps useful in describing natural ecosystems, as one does not need to know the precise location of the transition point and the specific temporal changes. Hence, while the observed KZ lag shares similarities with the concept of ‘extinction debt’ (Tilman et al., 1994), it exhibits unique characteristics associated with the critical slowing down at the transition. The presented KZ analysis could be relevant to understand the diversity dynamics in ecosystems undergoing climate change. In tropical soils, changes in conditions such as phosphorous bio-availability (which is sensitive to climatic changes that regulate soil redox) may cross a transition point and lead to a similar lag in response.

### Limitations and future directions

One possible criticism of our work is that the model we studied may not be general enough to capture the complexity of real ecosystems. Consumer-resource models are widely used in microbial ecology and provide a valuable tool for understanding the interactions between microorganisms and their environment. Additionally, since we focused on the scaling behavior near the transition point, we expect our results to apply to a broad universality class of systems that behave similarly near transitions. Therefore, our findings may apply beyond the minimal model studied here. We expect that there exist both relevant and irrelevant perturbations to our model that could affect the critical scaling we observed. One interesting direction involves considering the distribution of species in phenotype space, which here we take as uniform, but in reality may be quite different (Rubin et al., 2023). We leave a deeper investigation of these questions to future work.

A second criticism is that natural systems may not exhibit enough orders of magnitude for the ξ divergence to be observable, and therefore the slowdown may not be apparent. We should not conflate our inability to measure species abundances accurately with the fact that many natural systems exhibit a broad range of abundances spanning multiple orders of magnitude. Particularly, microbes are generally found in large numbers, indicating that real ecosystems—with thousands of species and abundance spanning several orders of magnitude—can, in principle, exhibit significant slowing down.

To test our KZ predictions, experiments in controlled laboratory conditions would be most beneficial. The primary aim would be to test the predicted lag in response to environmental change. An ideal experiment might commence with inocula sourced from fresh tropical rainforest soil samples, ensuring a high species richness. Initially, one must confirm in the serial-dilution lab setting, the attainment of a diverse community steady state which responds to changes in phosphorus. This would involve numerous serial-dilution steps, with subsequent ribosomal 16S amplicon sequencing to verify the establishment of a diverse steady state. Next, one would establish the existence of a low-diversity state by altering bio-available phosphorus concentrations through changing the environmental conditions. This could be done by altering soil oxygen levels and redox states, which influence microbial activity through changes in phosphorus availability and enzymatic activity (Gross et al., 2020). 16S sequencing would again determine the steady-state microbial composition.

Once the two extreme states—diverse and non-diverse—are established, it is not necessary to pinpoint the precise location of the transition point. Rather, one would then carry out several serial-dilution experiments at intermediate values between these two extremes, measuring the steady-state diversity. These experiments would determine the system’s steady-state behavior, mirroring a curve similar to Fig. 4a. Next, starting from the diverse state, conditions would be gradually altered from batch to batch, measuring each batch without allowing the system to reach steady state. Our theory predicts a lagged response, leading to behavior analogous to Fig. 4c. By contrasting the steady-state and gradual measurements, the data may reveal the presence of a lag.

Should 16S sequencing prove too costly or complex, in principle, one could measure any ecosystem-wide characteristic. Critical scaling theory suggests that as the entire ecosystem slows near the transition, these slow dynamics would manifest in any system-wide quantity. Thus, other potential measurable system characteristics include its carbon use efficiency (CUE), microbial activity, CO_2_ production, etc. We expect a similar lag, due to critical slowing down, when observing these quantities.

## CONCLUSIONS

There is considerable interest in quantifying ecosystem resilience and identifying early-warning signals of collapse (Ingrisch & Bahn, 2018; Bury et al., 2021). We build upon these foundations by examining critical slowing down in serial-dilution consumer-resource models and highlighting a potential lag, with significant implications. If this lag exists in real ecosystems, current conservation efforts may overlook ongoing extinction events. However, this lag also harbors a glimmer of hope: ecosystems that have been driven beyond their loss-of-diversity threshold can still be rescued if conditions are rapidly rectified. Therefore, we are eager to find out if the theory proposed here applies to real systems. However, in light of the alarming loss of diversity in natural ecosystems in recent decades, we remain hopeful that our theory’s most dire predictions will not be realized.

## ACKNOWLEDGMENTS

We thank Guy Bunin for comments about the scaling, and Jaime G. Lopez for critically reading the manuscript.

## AUTHOR CONTRIBUTIONS

AE conceived the theoretical idea and together with AG outlined its relevance to rainforest soils. AB, MGH, and AE wrote numerical simulations and made analytical calculations. All authors wrote the manuscript.

## DATA AVAILABILITY STATEMENT

All code and data used in this manuscript can be found at https://github.com/AmirErez/LossOfDiversity.

## APPENDIX

This section provides further details on the methods used in the manuscript “Diversity loss in microbial ecosystems under gradual environmental changes”.

### A EXTRACTING THE RELAXATION LENGTH FROM SIMULATION TIMESERIES

To extract the relaxation length from the convergence of the Shannon entropy to steady state, we faced a challenge. The theoretical way to do this is by fitting an exponent to *S*_*b*_–*S*_∞_ = *Be*^−*Cb*^ with *S*_∞_ the steady state Shannon entropy. Since only close to steady state we can expect the convergence to be dominated by a single scale, we considered convergence to within 1/100 distance from its steady state value. Nevertheless, the exact value of *S*_∞_ is unknown for *d* > 0, though we know it is small since *d* → 0. Small numerical differences mean that we cannot simply use the value of *S* at the final batch, since it may be slightly different from the true value. When fitting exponential decay, this slight difference can become a significant hurdle. Therefore, we needed to fit the form *S*_*b*_ = *A* + *Be*^*Cb*^. Approaching this problem using nonlinear regression proves difficult. Rather, an elegant solution by integration comes from Jacquelin (Jacquelin, 2009; Lecca, Lecca, Maestri, & Scarpa, 2021):

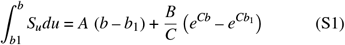

Replacing 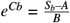 gives,

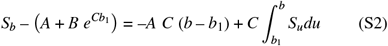

One may then numerically approximate the integral. Thereafter, the approximate solution and the exact solution are not the same, and minimizing the sum of the squared residuals between them results in a linear regression with respect to the coefficients *A, B*, which then leads to a second regression for *C*. For further details, see Jacquelin (Jacquelin, 2009) and https://scikitguess.readthedocs.io/en/latest/appendices/reei/translation.html. Together with this manuscript, we provide the function *shifted_exponential*.*m* which contains a MATLAB implementation of this method.

Though excellent, this fitting procedure is not perfect. Rather, we noticed that occasionally, the fit would converge to a solution that differs significantly from the data. Therefore, we removed outliers (implemented as *find_outliers*.*m*) that are at least 1.2-fold larger or smaller than their immediate neighborhood. This procedure removed about 1% of the data and filtered the failed fits almost perfectly.

**FIGURE S1.**
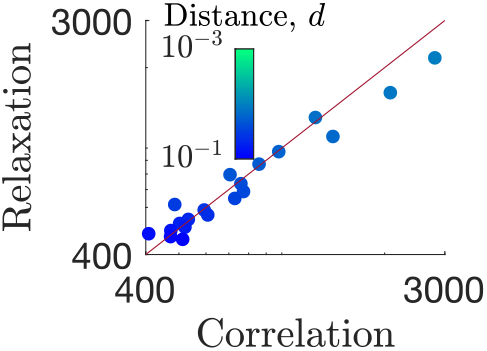
Batch-to-batch correlations in a stochastic system (*x*-axis, Gillespie simulation according to Table. S2) confirm that they are the same as the relaxation length of the equivalent deterministic system (*y*-axis, numerical solutions of Eq. 1). Line—equal axes. *K*=1, *c*_0_=10^2^, *ρ*_0_=2×10^4^, *α*=0.9.

### B STOCHASTIC SIMULATION

We used a stochastic simulation to show the equivalence of the correlation time of the stochastic system and the relaxation of the deterministic system. This simulation uses Gillespie’s algorithm where the birth and death reactions are with rates in accordance with their nutrient consumption rate. Note that in this case the species abundances are natural numbers and the nutrient abundances are real numbers. We simulated the reactions shown in Table S2 until all nutrients {*c*_*i*_} were depleted. For each “birth” reaction, (*ρ*_*σ*_ increases by 1), *c*_*i*_ decreases by *j*_*σ*,i_/Σ_*i*_*j*_*σ,i*_.

The normal serial dilution procedure is carried out and the system is allowed to reach steady state. Then, to extract the correlation, we use the method of batch means (Thompson, 2010). (Note: here, a single batch in the serial-dilution sense is collected to a batch of batches, an unfortunate jargon clash from two different fields). The idea is to divide a simulation trajectory of length *X* into batches of length *ξ*_*b*_. In the limit *X* ≫ *ξ*_*b*_ ≫ *ξ*, the correlation can be estimated by

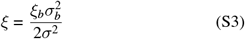

where 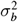 is the variance of the means of the batches and *σ* is the total variance (Thompson, 2010). We have used this method previously with good results (Byrd et al., 2019). A comparison between the two scales (deterministic relaxation, stochastic correlation) is shown in Fig. S1.

### C ANALYTIC FORM OF THE RELAXATION LENGTH

We present a theoretical derivation to calculate the *d* → 0 scaling in the chemostat limit, *c*_0_ ≪ *K*. We will assume symmetric strategies of two species and two nutrients, with *α*_1_ = (*α*, 1 – *α*) and *α*_2_ = (1 – *α, α*). We define 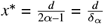with *δ*_*α*_ = 2*α*– 1.

**TABLE S1.** Annotation glossary.

| Symbol | Description |
| --- | --- |
| $t$ | Time measured from the beginning of a batch |
| $p$ | Number of nutrients |
| $m$ | Number of distinct species of the initial inoculum, the maximal number of species in the system |
| $m_e$ | Effective number of species at steady state |
| $i$ | (1...$p$) Latin index enumerating nutrients |
| $c_0$ | $\sum_{i=1}^p c_i(0)$; total nutrient concentration at time $t = 0$ |
| $c_i(t)$ | Time dependent concentration of nutrient $i$ |
| $P$ | Relative concentration of the first nutrient at time 0, $c_1(0)/c_0$ |
| $P_c$ | The value of $P$ at the loss-of-diversity transition point |
| $d$ | Equivalent to $P_c - P$, the distance between nutrient composition $c_1(0)$ and loss-of-diversity point |
| $K$ | Monod half-velocity constant |
| $\sigma, \sigma', \dots$ | (1...$m$) Greek indices enumerating species |
| $\rho_\sigma(t)$ | Species $\sigma$ biomass density at time $t$ since the start of a batch |
| $\vec{\alpha}_\sigma$ | $(\alpha_{\sigma,1}, \dots, \alpha_{\sigma,p})$; metabolic enzyme allocation strategy for species $\sigma$ |
| $j_{\sigma,i}$ | Nutrient $i$ consumption rate by species $\sigma$ |

**TABLE S2.** Gillespie reactions for resource competition dynamics

| Name | Reaction | Rate |
| --- | --- | --- |
| Birth | $\sigma \rightarrow 2\sigma$ | $\rho_\sigma \sum_i j_{\sigma,i}$ |
| Time | $t \rightarrow t + \Delta t$ | $\Delta t = -\ln(U(0, 1))/\sum_{\sigma,i} \rho_\sigma j_{\sigma,i}$ |

The steady state solution is 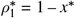 and 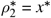. Let

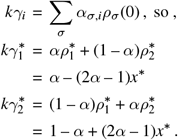

For batch *b* let *x*^(*b*)^ = *x*^*^ + Δ*x*^(*b*)^, with Δ*x*^(*b*)^ a small perturbation to the system at batch *b*. We have,

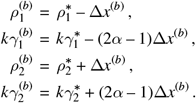

For *c*_0_ ≪ *K* we have *I* ≈ *ϕ* (Erez, Lopez, et al., 2020), giving,

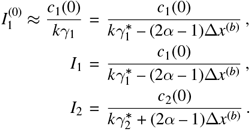

When the batch is completed, at time *t*_*f*_,

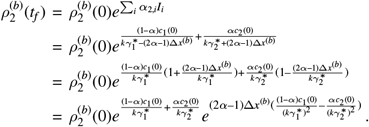

But 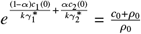, and so,

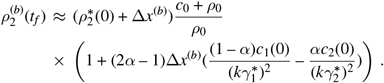

On the other hand, we have,

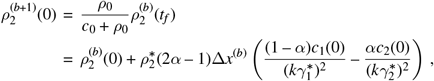

but, 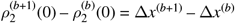. And so,

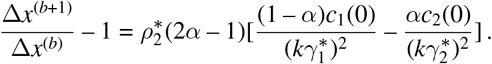

We consider that 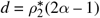 and we define *R* as the factor,

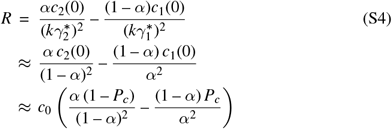

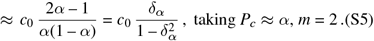

Given that the steady-state values 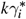 also depend on *d*, keeping the leading order in *d* gives 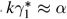 and 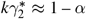, which gives Eq. 3. Furthermore, although *c*_1_(0) depends on *d*, it is by a small correction, 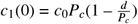 and therefore the *d* order may be dropped.

By setting 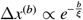, we get,

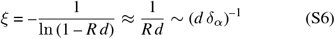

Therefore, when considering *ξ* ∼ *d*^−*νz*^ we have *νz* = 1. We may change 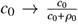 for a more general dependence that recognizes that multiple batches are needed to make any change when *c*_0_ < *ρ*_0_. We see the *ξ* ∼ |*dδ*_*α*_|^−1^ in both Fig. 3 for *c*_0_ = 10 with further plots shown in Fig. S2. The same analysis for *c*_0_ = 10^3^ is shown in Fig. S3.

### D SHANNON DIVERSITY NEAR THE TRANSITION POINT

To derive the dependence of the Shannon diversity on distance near transition point, we first establish the applicability of an Ansatz expression for the steady state {*ρ*_*σ*_}, and then use this Ansatz to calculate the Shannon diversity.

Let a system with *m* species have strategies equally spaced between (*α*, 1 – *α*) and (1 – *α, α*), as in the results presented in Fig. 3 and Fig. S3. Let the initial inocula species abundances (of the first batch) be equal for all species. At steady state, the resulting species abundance distribution follows the Ansatz,

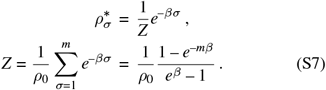

We do not currently have analytical arguments to prove this Ansatz, however, it is easy to demonstrate that it works well in practice. We attempted to fit all the {*ρ*_*σ*_(*t* = 0)} data used for Fig. 4. An example of the fit for *m* = 8 and several values of *d* is shown in Fig. S4a. We have carried out the fit on all the simulation output used for Fig. 4 and calculated the sum of squared errors (SSE) divided by the number of data points. The fit works exceptionally well in all cases (Fig. S4b).

By fitting the Ansatz for {*ρ*_*σ*_}, Eq. S7, to the data, we extract *β* for all *m, d* pairs. We then follow the same logic as for the large *d* limit: the maximal entropy of the system gives *e*^*S*^ ≡ *m*_*e*_ = *m* and therefore (*m*_*e*_ – 1) ∼ ℳ(*d*/*δ*_*α*_) is a good candidate. Plotting *β* vs. 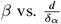 suggests a phenomenological relation *β* = −*q* In 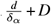,at small 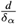,, see Fig. S4c. We insert the expression for *β* to express the entropy as a function of 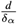, in particular *e*^*s*^−1.

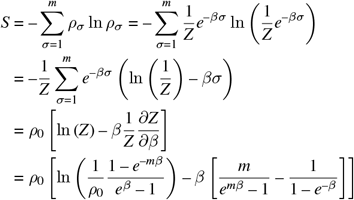

Substituting 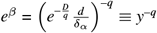 and taking the limit 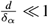, we get,

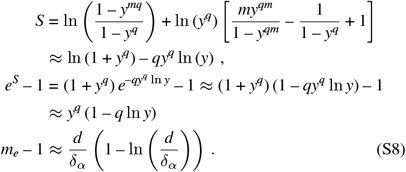

This relation between *m*_*e*_ and 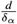 at small 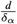 is precisely the relation shown in Fig. 4b. We note that it has a similar, though distinct slope on the scaling collapse curve, than the slope 1 when 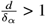.

**FIGURE S2.**
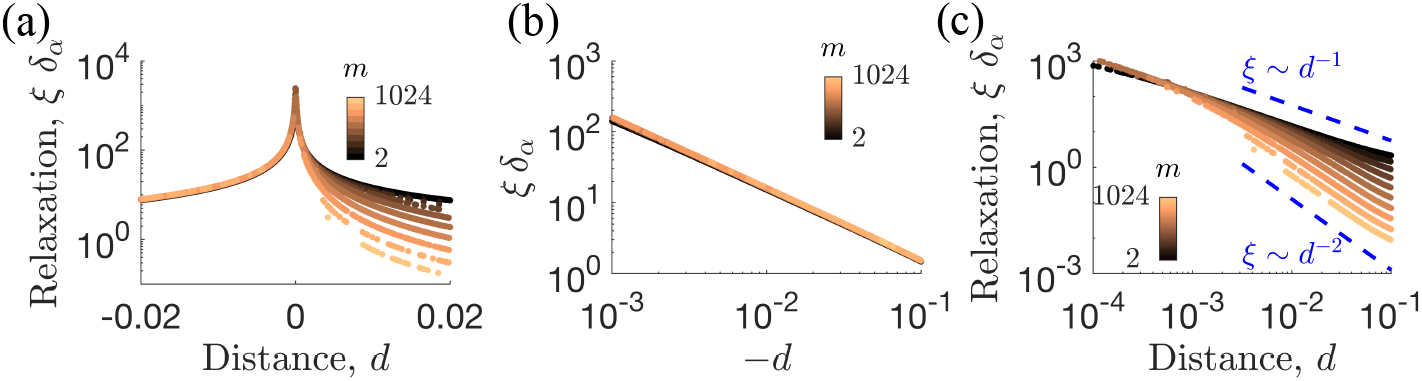
Alternative scaling presentations of the relaxation length from Fig. 3 in the main text. (a) The equivalent of Fig. 3a but on a logarithmic scale on the *y*-axis shows clearly the different scaling for *d* < 0 and *d* > 0 regimes. (b) When scaling *ξδ*_*α*_ ∼ (–*d*^−1^) we see a full scaling collapse of the *d* < 0 branch presented in Fig. 3b. (c) The *d* > 0 branch, shown in Fig. 3c, viewed with *ξδ*_*α*_ on the *y*-axis. At small *d* we see the same scaling for all *m* values, but a with crossover to *m*-dependent behavior at larger *d*.

**FIGURE S3.**
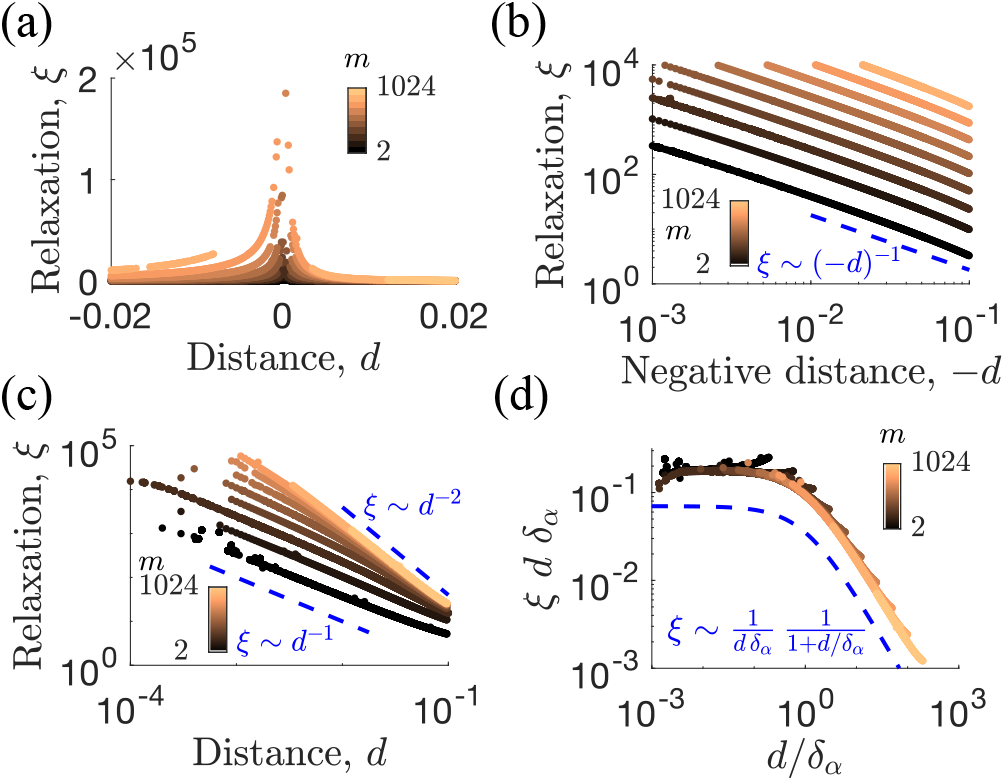
Numerical simulations of the number of batches required for relaxation, *ξ*, as a function of the distance to the transition point, *d*, with *d* > 0 marking the region where the nutrient composition supports diverse steady states. This figure is identical to Fig. 3 except the data were simulated with *c*_0_ = 10^3^ instead of 10^1^.

**FIGURE S4.**
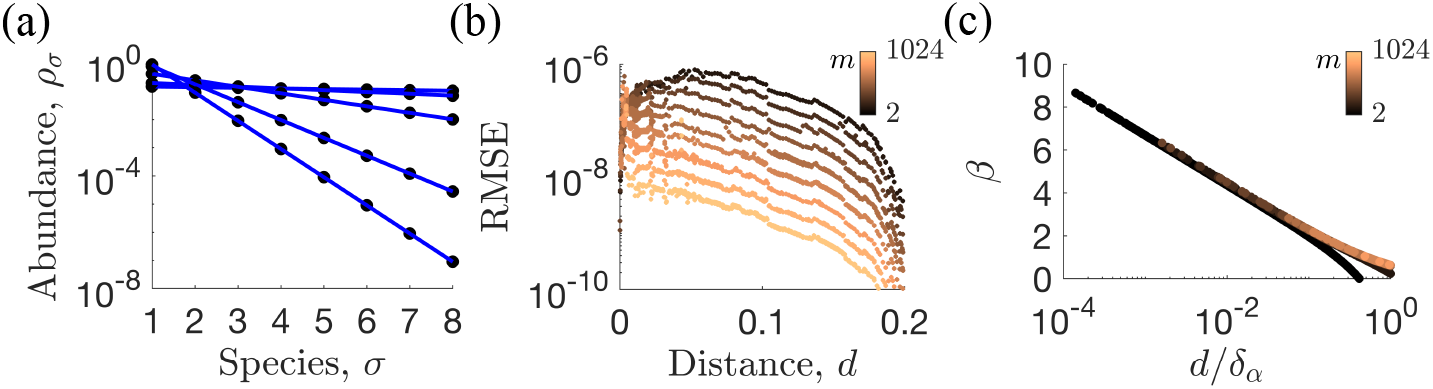
Ansatz for the species abundance distribution according to 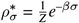. (a) Example of the fit of the species abundances (numerical results, dots) for various distances from the transition point (different curves). The fit is according to the Ansatz in Eq. S7 (lines). (b) Root-mean-squared-error (RMSE) of the fit, carried out on all the data used in Fig. 4. As is apparent, the fit works exceptionally well in all cases. (c) For 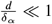, the decay factor *β* collapses for all species, and is linear in log 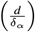. Fitting this expression allows us to express the entropy of the system as a function of 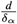.

